# Orchestrating multi-state QTL analysis with Bioconductor

**DOI:** 10.1101/2025.03.28.645834

**Authors:** Christina B. Del Azodi, Amelia M. Dunstone, Davis J. McCarthy

## Abstract

**Motivation:** The scope of many Quantitative Trait Loci (QTL) mapping studies has increased to include different cellular and environmental states. However, drawing biologically relevant conclusions from the large, high-dimensional data that come from multi-state QTL mapping studies is not straightforward.

**Results:** To address this problem, we introduce two R packages, *QTLExperiment* and *multistateQTL*. The *QTLExperiment* package provides a robust container for storing and manipulating QTL summary statistics and associated metadata. Building upon existing Bioconductor infrastructure and conventions, this object class is consistent, user-friendly, and well-documented. The *multistateQTL* package introduces tools to facilitate the analysis of multi-state QTL data stored in a *QTLExperiment* container. This package provides methods for statistical analysis, quantification of sharing, classification of multi-state QTL associations, visualization of the data, and more. It also provides flexible methods for simulating multi-state QTL summary statistics with user-defined properties.

**Availability and implementation:** The packages *QTLExperiment* and *multistateQTL* are available on Bioconductor (https://www.bioconductor.org/packages/QTLExperiment and https://bioconductor.org/packages/multistateQTL).

## Background

Quantitative Trait Loci (QTL) mapping studies test for associations between genetic variants such as single nucleotide polymorphisms (SNPs) and changes in a molecular trait like gene expression (eQTL) or chromatin accessibility (caQTL). These associations provide insights into the regulatory basis of traits. As sequencing technologies advance, the size and scope of QTL mapping studies continue to grow. A major area of growth has been in expanding the cellular contexts in which QTL are assessed, for example in different tissues (e.g., [1, 2]), in specific cell-types (e.g., [3]), or during specific developmental stages (e.g., [4]). QTL studies have also expanded to consider environmental (e.g., [5, 6]) and population (e.g., [7, 8]) contexts. We refer to studies that test for QTL associations across more than one of these contexts as multi-state QTL.

There are many established frameworks for performing QTL mapping and a growing number of tools for the joint analysis of multi-state QTL [9–11]. The output from these tools is a set of summary statistics for each association in each state, including the estimated effect size, the variance or standard error around that estimate, and often a test-statistic describing the significance of the association (e.g., false discovery rate, FDR, corrected p-values, local false sign rates, LFSRs [12]). This big, high-dimensional data presents a number of challenges, including how to describe global patterns of sharing and how to identify groups of related associations, and specific associations that may be of biological interest. To help facilitate these goals, here we introduce two packages:

1. *QTLExperiment*: a container for storing and manipulating QTL summary statistics and associated metadata; and
2. *multistateQTL*: a toolkit for simulating, analyzing, and visualizing multi-state QTL summary statistics.

The *QTLExperiment* container is designed to store important metadata along with the summary statistics to allow for robust data merging and subsetting, with checks in place to reduce the potential for labeling mistakes. The *multistateQTL* package builds on *QTLExperiment* to aid users in the analyses of complex QTL datasets.

These packages are intended to help the community: feedback or feature requests are appreciated and can be submitted on GitLab (https://gitlab.svi.edu.au/biocellgen-public/qtlexperiment/-/issues; https://gitlab.svi.edu.au/biocellgen-public/multistateqtl/-/issues)

## 2 Methods, data, and implementation

### 2.1 Data usage

The empirical results presented in this paper were generated using expression QTL (eQTL) summary statistics from the Genotype-Tissue Expression (GTEx) Project (version 8; [2]) for the ten tissues with the largest sample sizes for eQTL mapping. The eQTL tests were filtered to include only those on chromosome 1 that were also available in all 10 tissues. These filtered data were also used to estimate parameters needed for the simulations (described below), and to set the default parameters available in *multistateQTL*.

### 2.2 Implementation

The *QTLExperiment* and *multistateQTL* packages are open-source R packages. Both packages take advantage of cutting-edge R programming tools to ensure speed and memory efficiency. The sumstats2qtle() function can load over 20 million beta, error, and p-value summary statistics from 10 states in 10 minutes. Once loaded into a *QTLExperiment* object, calling significant tests takes 4 seconds, filtering to keep only the top hits per feature takes 25 seconds, and calculating and plotting pairwise sharing on those top hits takes less than a second.

*QTLExperiment* builds on *SummarizedExperiment* from Bioconductor [13]. Both packages utilize existing CRAN and Bioconductor packages for efficient data manipulation and plotting. For example, *vroom* [14] is used for loading data with parallel processing when multiple cores are available and *collapse* [15], which uses C/C++ based functions, is used for intensive data transformations. The plotting functions use *ggplot2* [16] and *ComplexHeatmap* [17].

## 3 Results

### 3.1 The *QTLExperiment* container

The *QTLExperiment* container is a lightweight, S4 class object for storing summary statistics and metadata from QTL mapping workflows. It extends the *SummarizedExperiment* class and is organized such that rows represent QTL associations between a feature and a variant, columns represent states, and assays contain various summary statistics (**Figure 1**).

**Figure 1.**
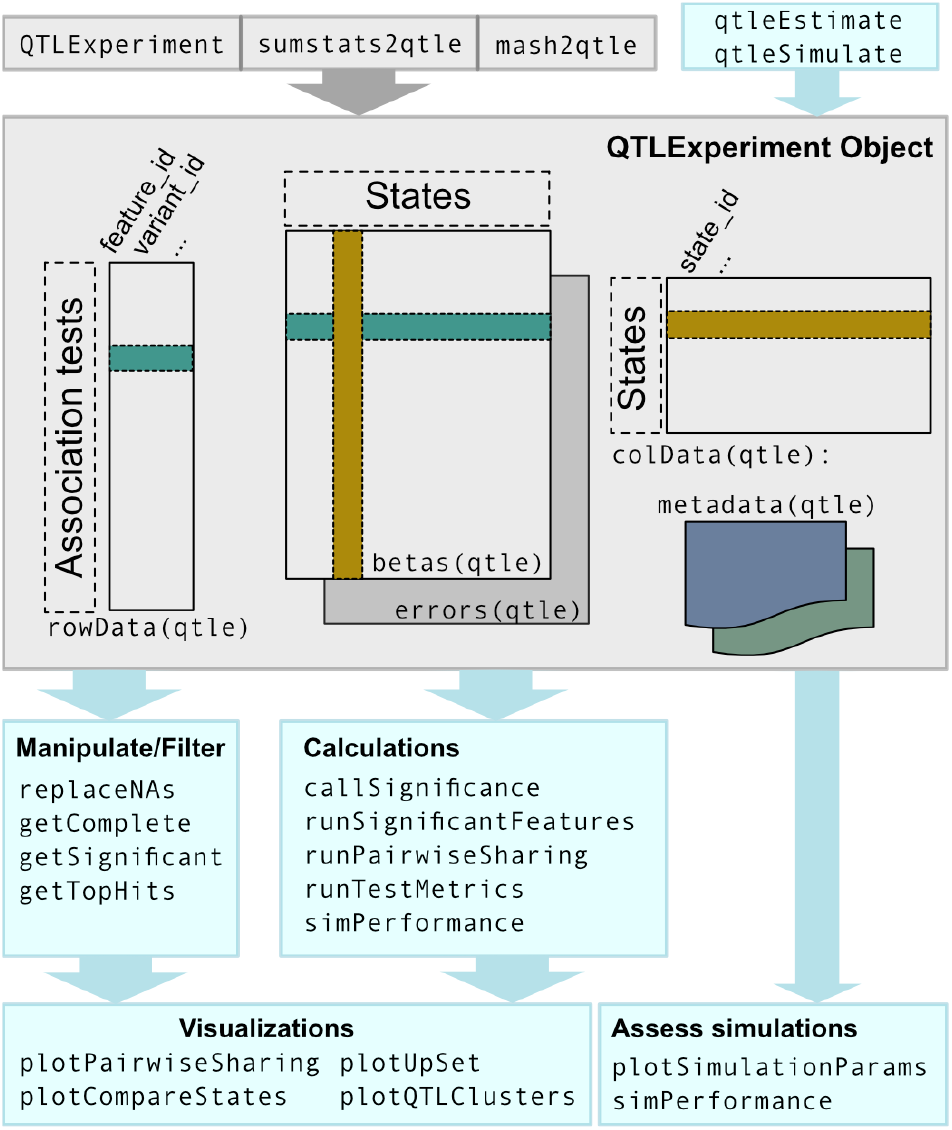
Overview of the *QTLExperiment* object class and available methods. Methods available in the *QTLExperiment* package are shaded in gray, and those implemented in the *multistateQTL* toolkit are shown in the blue boxes.

The *QTLExperiment* class inherits the slots rowData, colData and assays from the parent class *SummarizedExperiment*, with some additional requirements on the slots enabling the storage of multi-state QTL data. The class has two required assays: “betas”, for storing the estimated QTL effect sizes; and “errors”, for the effect sizes (typically standard errors) around each estimate. Any number of additional assays (e.g., test-statistics, known/simulated betas) can be stored. The rowData stores metadata about each QTL association. The two required rowData columns are feature_id and variant_id, which store the feature names (e.g., gene names) and the variant information (i.e., SNP names), respectively. Each row is required to have a unique combination of these two components. The colData stores metadata related to the states and has one required column, state_id. The state ID names must be unique. Additional metadata that needs to be stored with the instance, but does not directly relate to associations or states, for example covariance matrices used for QTL mapping, can be stored in the metadata list.

The *QTLExperiment* object stores data in a robust fashion, keeping metadata organized and special fields protected when the object is manipulated. Due to the importance of the feature_id, variant_id, and state_id information to the object, there are several protective measures in place to maintain the validity of this data. When updating these fields, the methods in *QTLExperiment* ensure that the new values provided are sensible (e.g., new values should have a one-to-one mapping with the old values). In addition, overwriting the colData or rowData of a *QTLExperiment* object will not delete data from the protected fields. The package also includes specific functions for the protected fields, namely state_id(), feature_id() and variant_id(), which provide a shorthand for viewing or changing these fields in the row or column metadata. The package includes methods for subsetting the object, as well as for binding rows and columns. Together these features make the *QTLExperiment* object user-friendly while also preserving important properties about the data.

Finally, the package contains a suite of functions to convert user data into a *QTLExperiment* object. This includes a function to read in QTL summary statistics from separate files for each state. This function is flexible, allowing the user to provide QTL summary statistics as output from a range of common QTL mapping software tools or data resources (e.g., *FastQTL* [18], *LIMIX* [19], GTEx, etc.), by specifying where key data is stored in those files. It also includes specific functions to read in joint multi-state QTL summary statistics from common intermediate tools (e.g., *mashr* [10]).

### 3.2 The *multistateQTL* toolkit

The *multistateQTL* package provides a wealth of tools to assist with analyzing multi-state QTL summary statistics. It includes functions for filtering and imputing missing assay data, calling significance, calculating pairwise sharing across states, selecting top QTL hits, and categorizing QTL by their pattern across states (**Figure 1**). Here we demonstrate these tools using all QTL tested on chromosome 1 across 10 GTEx tissues.

#### Dealing with missing data

Some common downstream analysis steps cannot handle missing data in the *QTLExperiment* object, specifically in the betas and errors assays. Thus, we developed methods to filter and/or impute missing data. The filtering function getComplete() allows the user to specify a minimum number of states with complete data for the QTL to be retained. The imputation function replaceNAs() allows the user to specify for each assay how the NAs should be replaced. This flexible approach could be used to assign NA effect sizes a value of zero, NA p-values a value of one, and NA standard errors the row (i.e., test) median values. We used the getComplete() function to remove ∼8.4 million QTL tests that had missing data in one or more of the 10 GTEx tissues, leaving ∼11.6 million QTL for the downstream analysis.

#### Calling significant associations

QTL tests can be called as significant using either a one-step or two-step threshold approach using the callSignificance() function. The one-step threshold approach simply calls QTL as significant in a given state if the test statistic specified by the user is below a given threshold (e.g., LFSR < 0.05). The two-step threshold approach allows for a more lenient threshold for calling significance in additional states if the QTL is significant at the stricter threshold in at least one state. This more flexible approach can help minimize the impact of differential power across states, which commonly arises when different states (e.g., tissue, cell type) have different sample sizes. For our GTEx analysis, we applied the two-step threshold approach (p-values < 0.1 for the first threshold, p-values < 0.2 for the second).

#### Summarizing results by feature

In single-state QTL mapping, results are often filtered to only include the top QTL per feature (e.g., the top eSNP per gene). However, in multi-state QTL analysis this filtering is complicated because the top QTL may be different in different states for reasons either biological (e.g., lead-switching) or technical (e.g., linkage disequilibrium, LD, between eSNPs). This complication can be avoided by summarizing multi-state QTL results on the feature level. The runSignificantFeature() function adds a summary of the features with significant QTL in each state to the metadata of the *QTLExperiment* object for downstream analysis. In addition, the getTopHits() function can be used to select the top QTL per feature either globally, where the test with the lowest test statistic per feature across all states is selected, or at the state-level, where the tests with the lowest test statistic per feature per state (in states where the feature is significant) are selected. For our GTEx analysis, filtering to keep the top hits per gene per state resulted in 14,761 QTL used for the downstream analysis.

#### Calculating pairwise sharing

To quantify the degree of pairwise QTL sharing between states, we implement the approach used in *mashr* [10] in our runPairwiseSharing() function. Using this approach, a QTL is considered shared if it is significant in either state and the effect sizes are within a factor of each other. As per *mashr*, the default factor is set to 0.5, however users can increase or decrease this factor to require higher or lower similarity between effect sizes, respectively. At the extreme ends, setting the factor to zero tests for a shared direction regardless of magnitude, while setting the function parameter to the absolute value (FUN=abs) tests for shared magnitude regardless of direction. For our GTEx analysis, we calculated the pairwise sharing on the top QTL hits using the default approach and then plotted these results using the plotPairwiseSharing() function. Without any multi-tissue integration being applied, we found between 49% and 61% of QTL were shared between tissue pairs, with similar tissues (e.g., “skin -sun exposed” and “skin -not sun exposed”) having the most sharing (**Figure 2a**).

**Figure 2.**
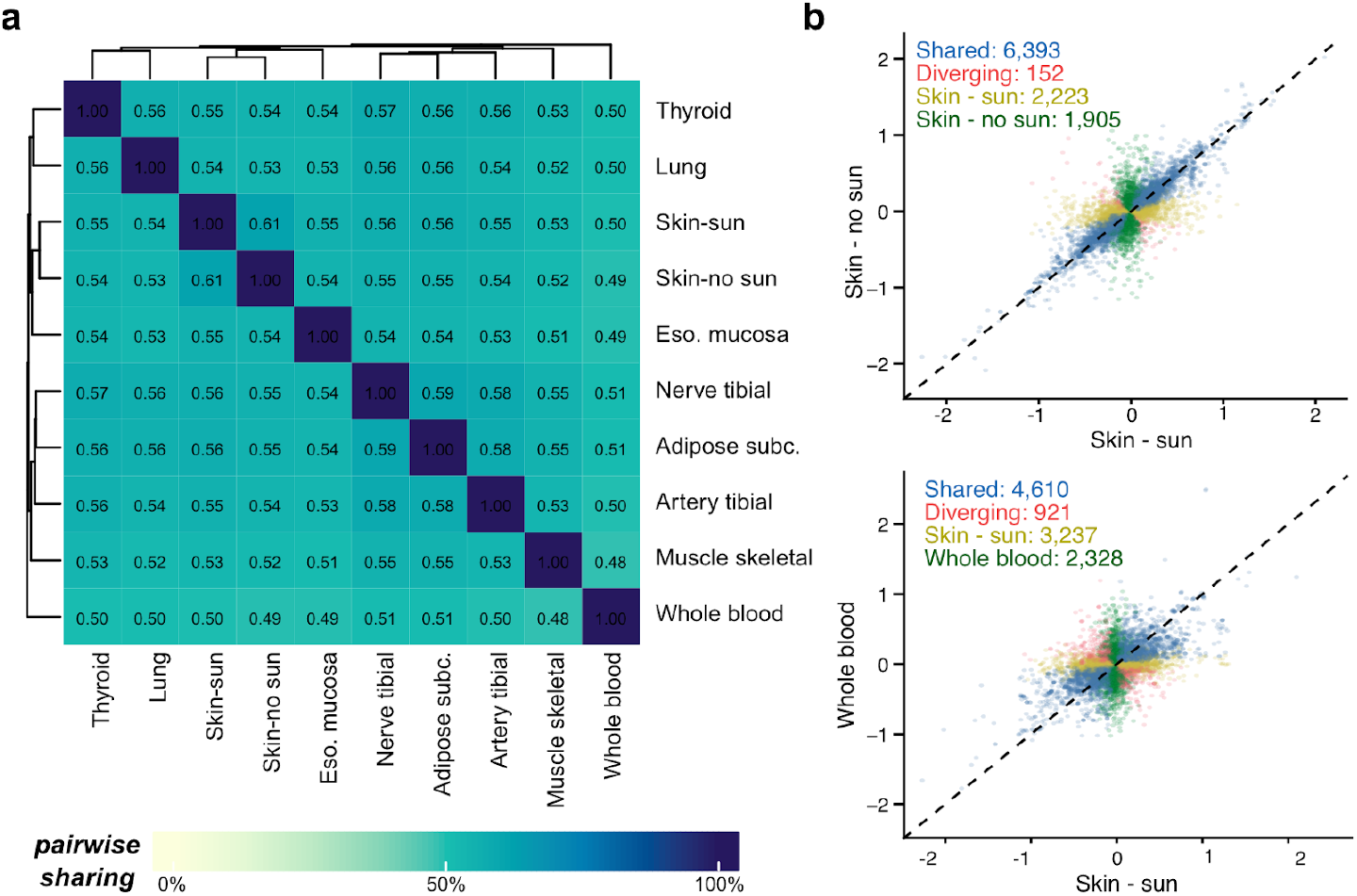
Global summary of GTEx chromosome 1 multi-tissue eQTL. **(a)** Pairwise sharing of eQTL across tissues, tissues are sorted by hierarchical clustering. **(b)** Comparison of estimated eQTL effect sizes between two tissues of interest. eQTL that are shared between tissues are shown in blue, those with diverging effects are in red, those only significant in the tissue on the x-axis are in yellow, and those only significant in the tissue on the y-axis are in green.

#### Categorizing multi-state QTL

We introduce standard definitions for categories of multi-state QTL based on their pattern of sharing and effect direction across states. The broad categories describe QTL as “global” if they are significant across all states, “multi-state” if they are significant across only a subset of states, and “unique” if they are specific to a single state. Global and multi-state QTL are further categorized by the pattern of the effect sizes in states where the QTL is significant. If the QTL has the same effect direction in all significant states, it is considered “shared”. If the QTL has significant effects in the opposite direction in at least one state, it is considered “diverging”. After calling significant tests, the runTestMetrics() function can be applied to categorize QTL using these definitions. It also calculates the variance in effect sizes across states where the QTL is significant and reports the number of significant states for each QTL. For our GTEx analysis, after applying runTestMetrics(), we visualized and quantified these multi-state QTL categories in two similar (“skin -sun exposed” and “skin -not sun exposed”) and two very different (“skin -sun” and whole blood) tissues using the plotCompareStates() function (**Figure 2b**).

### 3.3 Simulation tools

The *multistateQTL* toolkit provides a framework to simulate realistic multi-state QTL summary statistics. The simulation framework consists of two parts: first, the estimation of key parameters from empirical QTL summary statistics; and second, the sampling of QTL summary statistics from these parameters given the user defined structure.

The default parameters available in *multistateQTL* were estimated from eSNP:eGene pairs tested on chromosome 1 in GTEx (v8) for the 10 tissues with the largest sample sizes (**Figure S1**). Users can estimate parameters from their own empirical summary statistic data to simulate datasets with desired characteristics. The shape and rate parameters define gamma distributions for the following components: the mean absolute value of effect sizes across states for significant QTL tests (betas.sig), the coefficient of variation (cv) of significant effect sizes across states (cv.sig), the absolute value of effect sizes across all null tests (betas.null), and the cv of null effect sizes across all null tests (cv.null).

The simulation tool allows for the simulation of four types of associations: (1) Global, where the simulated effect size is approximately equal across all states; (2) Unique, where the association is only significant in one state; (3) Multi-state, where the association is significant in a subset of states (i.e., state-groups), and (4) Null, where the association has no significant effects in any state. First, each test is randomly assigned as one of the above types according to the proportions specified by the user. For multi-state QTL, each state is assigned to a state-group, either randomly or according to user defined groups, then each multi-state QTL is assigned randomly to one of the state-groups. For unique QTL, the QTL is randomly assigned to a single state.

Simulated mean effect sizes for all non-null QTL are sampled from *gamma(beta*.*sig*.*shape, beta*.*sig*.*rate)* distributions and are randomly assigned a positive or negative effect direction. Then for each state where that QTL is significant, an effect size is sampled from *N(mean effect size, σ)*, where *σ* is user defined (default=0.1). Effect sizes for null QTL are sampled from *gamma(beta*.*null*.*shape, beta*.*null*.*rate)* and are randomly assigned a positive or negative effect direction. Standard errors for each QTL for each state are simulated by sampling from *gamma(cv*.*sig*.*shape, cv*.*sig*.*rate)* or *gamma(cv*.*null*.*shape, cv*.*null*.*rate)* for significant and null QTL, respectively, and multiplying the sampled cv by the absolute value of the simulated beta for that QTL in that state.

Using this simulation framework, multi-state QTL summary statistics can be simulated to represent different real-world scenarios. For example, we can simulate a simple scenario where most QTL are shared (**Figure 3a**). Here we consider simulated QTL as shared if they are significant in either state and have simulated betas within a factor of 0.75. States were called as significant using the two-step threshold approach, with one state having an LFSR < 0.1 and the remaining states having an LFSR < 0.2. We can also specify different *σ* to be used for each state, allowing us to simulate a scenario where there is more variability in effect sizes in some states (e.g., S5: *σ = 1*, and *S6: σ = 2;* **Figure 3b**), which could happen if those states have fewer data or lower quality data available for QTL mapping. Finally, we can simulate scenarios with structure between states (e.g., two state-groups in **Figure 3c**), which would be expected, for example, if we were looking at QTL across different subgroups of tissues. Plotting the number of simulated QTL that are significant in different sets of states provides a more detailed look at the structure of QTL sharing within our simulation (**Figure 3d**). Finally, plotting and clustering the multi-state QTL by their sampled betas provides a detailed look at the properties of specific QTL (**Figure 3e**). Note that the heatmap and UpSet plotting functions allow for state and test level variables stored in the *QTLExperiment* colData and rowData, respectively, to be used to annotate the plots with contextual information (e.g., state-groups, number of significant QTL per state, and simulated mean beta; **Figure 3**).

**Figure 3.**
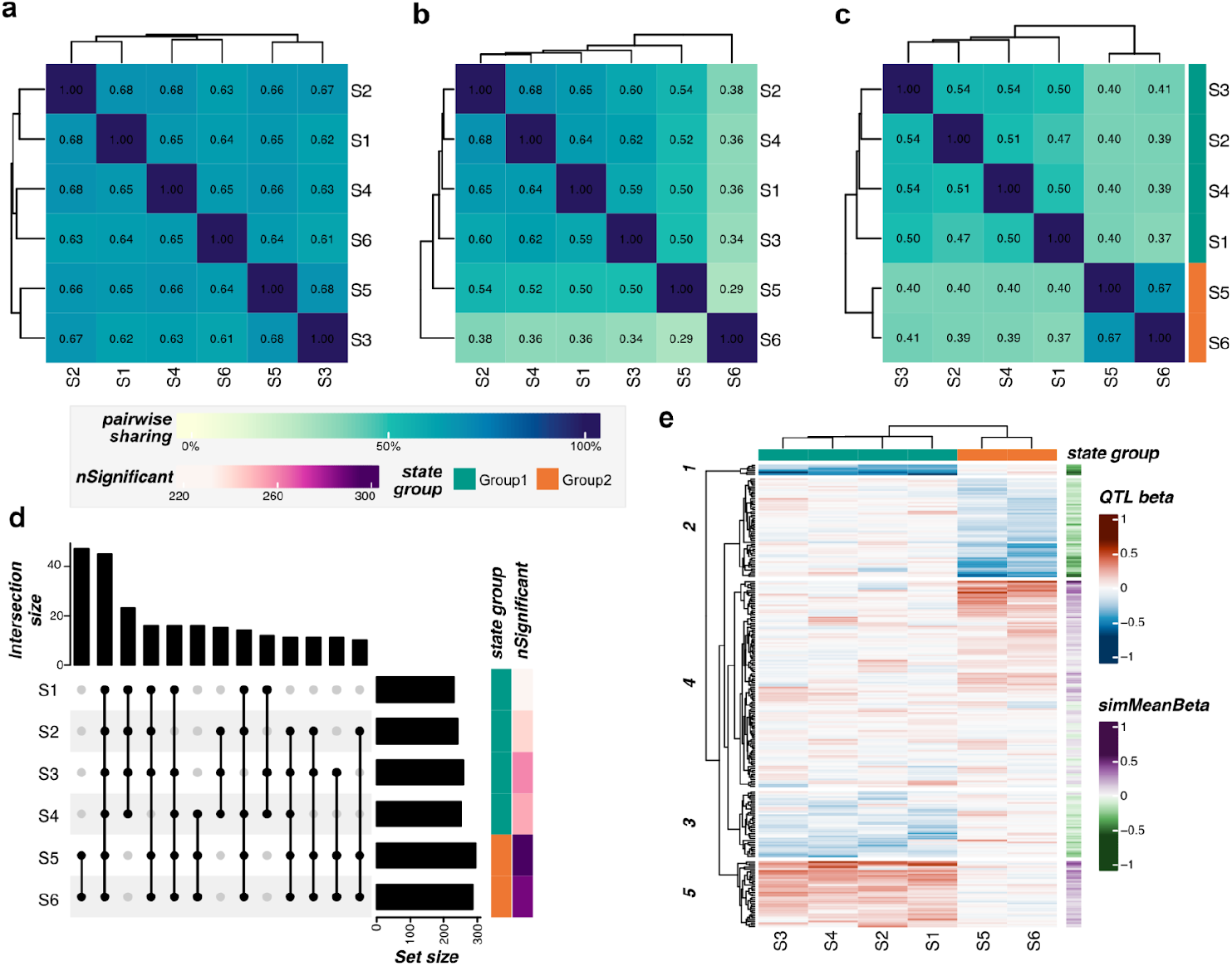
Simulated multi-state QTL. Pairwise sharing of 500 simulated QTL across 6 states for three simulation scenarios. (**a**) A simple scenario with 80% global and 20% state-specific QTL. (**b**) A scenario with different power for detecting QTL across states, with 80% global and 20% state-specific QTL, but where S1-S4 are simulated with betaSd=0.1 (default; high power), S5 with betaSd=1 (moderate power), and S6 with betaSd=2 (low power). (**c**) A scenario where there is state-level structure, created by simulating 50% of QTL with multi-state effects (i.e., effects assigned to only one state-group; here either S1-S4 or S5-S6), and 50% global QTL. (**d**) An UpSet plot showing the number of QTL (intersection size) for the simulation in **c** that are significant in sets of states shown by the black, connected dots. The number of QTL significant for each state is shown as the set size and in the nSignificant row annotation. (**e**) Sampled betas for multi-state QTL from simulation **c** clustered using hierarchical clustering with k=5. The QTL are annotated with the mean betas sampled for that QTL for significant states.

## 4 Conclusions

The *QTLExperiment* and *multistateQTL* R packages provide a user-friendly approach to loading, storing, manipulating, and analyzing multi-state QTL association summary statistics and provide tools for visualizing these data. Together these tools can be used to facilitate the analysis of high-dimensional multi-state QTL data to describe global patterns of gene regulation across states and highlight specific QTL and clusters of QTL that may be of particular biological interest. As the scope of QTL mapping studies grows to include more and higher resolution cellular and environmental contexts, software tools that facilitate the management and standardization of QTL data and workflows will become critical.

## Availability and requirements

The *QTLExperiment* and *multistateQTL* packages are implemented in R and are available through Bioconductor (https://www.bioconductor.org/packages/QTLExperiment and https://bioconductor.org/packages/multistateQTL). *QTLExperiment* is available in Bioconductor version 3.18 or higher, and *multistateQTL* requires version 3.19 or higher. It is recommended to use R version 4.4 or newer with these packages. Detailed vignettes are provided for each package, demonstrating example workflows and function usage. These software are licensed under the GNU General Public License v3.0. The code used to generate the results in this manuscript is available at https://biocellgen-public.svi.edu.au/WEEO_2022_multistateQTL/.

## Acknowledgements & Funding

We thank Jeffrey Pullin for testing and providing feedback on the software packages. This work was supported by the National Health and Medical Research Council of Australia [GNT116289 and GNT1195595 to D.J.M.], by the Baker Foundation [D.J.M. and C.B.D.], and through the St Vincent’s Institute of Medical Research Rising Star Fellowship [C.B.D.].

## Author contributions

C.B.D. and D.J.M. conceptualised the packages. C.B.D. developed the packages initially, including software code, supporting documentation, and vignettes. A.M.D. tested the software code and provided major updates to the source code and documentation. D.J.M. provided supervision for this project. All authors contributed to the final version of the manuscript.

