## Supplemental Figure 1 for "Orchestrating multi-state QTL analysis with Bioconductor"

### Supplemental Figures

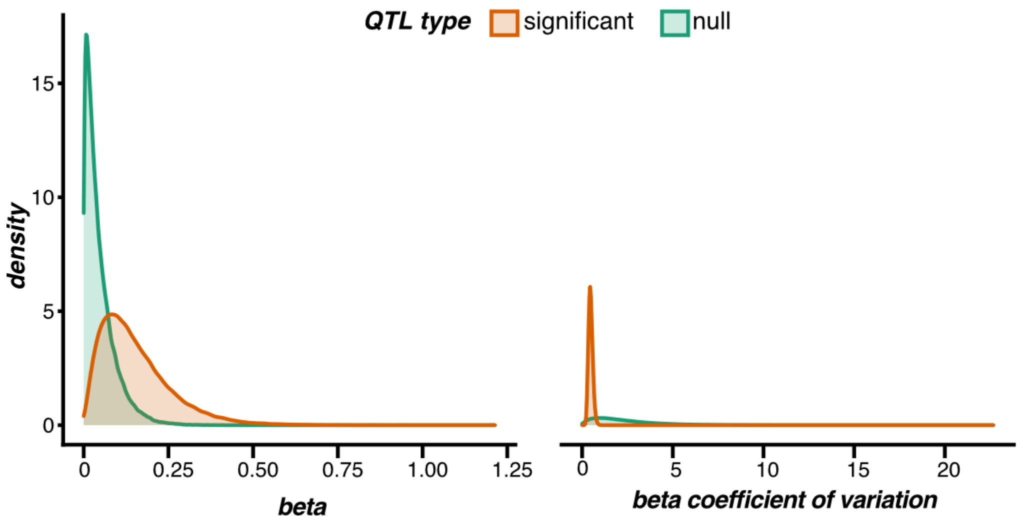


**Figure S1. Gamma distributions defined by the multi-state QTL parameters estimated from the GTEx data.**
